# Therapeutic dose prediction of α5-GABA receptor modulation from simulated EEG of depression severity

**DOI:** 10.1101/2024.05.15.594433

**Authors:** Alexandre Guet-McCreight, Frank Mazza, Thomas D. Prevot, Etienne Sibille, Etay Hay

## Abstract

Treatment for major depressive disorder (depression) often has partial efficacy and a large portion of patients are treatment resistant. Recent studies implicate reduced somatostatin (SST) interneuron inhibition in depression, and new pharmacology boosting this inhibition via positive allosteric modulators of α5-GABA_A_ receptors (α5-PAM) offers a promising effective treatment. However, testing the effect of α5-PAM on human brain activity is limited, meriting the use of detailed simulations. We utilized our previous detailed computational models of human depression microcircuits with reduced SST interneuron inhibition and α5-PAM effects, to simulate EEG of virtual subjects across depression severity and α5-PAM doses. We developed machine learning models that predicted optimal dose from EEG with high accuracy and recovered microcircuit activity and EEG. This study provides dose prediction models for α5-PAM administration based on EEG biomarkers of depression severity. Given limitations in doing the above in the living human brain, the results and tools we developed will facilitate translation of α5-PAM treatment to clinical use.

## Introduction

Major depressive disorder (depression) is a leading cause of disability, but the efficacy of current treatment methods is often partial or patients are treatment-resistant^1,2^, indicating disease mechanisms that the current methods do not address. In recent years, reduced cortical inhibition has been implicated as a mechanism in depression^3–8^, and new pharmacology that provided positive allosteric modulation of the α5 subunit of GABA_A_ (α5-GABA_A_) receptors (α5-PAM) in chronically stressed mice^9,10^ had antidepressant, anxiolytic, and pro-cognitive effects. However, testing the effect of the new drugs on human brain activity is currently limited. We have recently overcome these limitations by characterizing the effects of α5-PAM on cortical function and encephalography (EEG) measures of efficacy *in silico* using detailed models of human depression microcircuits^11^. This computational approach can further offer dose prediction tools via systematic characterization of simulated EEG effects across varying levels of depression severity and applied dose.

Due to their subunit selectivity, α5-PAM are optimized to selectively boost inhibition generated by somatostatin-expressing (SST) interneurons, which provide synaptic and extrasynaptic (tonic) inhibition to pyramidal (Pyr) neuron apical dendrites via α5-GABA_A_ receptors^12–15^. A loss of SST interneuron inhibition in depression is implicated by reduced SST expression in human patients postmortem^16,17^, increased anxiety- and depression-like symptoms in rodents with brain-wide SST interneuron silencing^18^, and SST interneuron transcriptome deregulations compared to other cell types following chronic stress in rodents^19^. This is further supported by the joint inhibitory effects and co-release of SST and GABA^20,21^. SST interneurons mediate lateral inhibition through disynaptic loops^22,23^ and maintain low Pyr neuron spike rates at baseline^24–26^. A reduced SST interneuron inhibition in depression would thus increase baseline cortical activity (noise) and impair signal-to-noise ratio (SNR) of cortical processing^8,27,28^.

Detailed biophysical models can capture key properties of EEG^29–31^, and thus provide powerful tools for linking cellular and circuit mechanisms to brain activity and clinically-relevant signals. We previously showed *in silico* that α5-PAM would recover cortical activity, function and EEG spectral profile in detailed models of human depression microcircuits with reduced SST interneuron inhibition, and we highlighted EEG biomarkers that can monitor α5-PAM efficacy^11^. However, the study utilized an average reduced SST interneuron inhibition as a model of depression microcircuits^28^ and EEG effects^30^, and did not consider varying depression severity level. A similar approach was used by detailed modeling studies that identified the effects of schizophrenia-related gene variants on neuronal cellular mechanisms and EEG features^32^. Other studies linked cellular and circuit mechanisms to features of the EEG response during task processing in health^33^ and in relation to schizophrenia biomarkers^34^.

EEG biomarkers have been used broadly in depression classification and treatment prediction. Particularly, elevated theta and alpha power are characteristic features in the frontal and parietal electrodes of depression patients^35–39^ as well as increased aperiodic activity^40^. We previously showed mechanistically, using detailed computational models, that increases in theta, alpha, and aperiodic power would result from reduced SST interneuron inhibition^30^. EEG power in different frequency bands also served to predict treatment response in depression^41^. In particular, theta and alpha power in depression patients have been used as indicators of treatment response^42– 46^, and similarly served as biomarkers in our previous simulated cortical microcircuit response to α5-PAM^11^.

In this work, we simulated virtual subjects with varying levels of depression severity in terms of reduced SST interneuron inhibition in detailed models of human cortical microcircuits. For each virtual subject we simulated a5-PAM dose response, and developed machine learning models to predict optimal dose from EEG biomarkers for recovering cortical microcircuit activity, function and EEG profile.

## Results

We simulated α5-PAM dose effects on detailed models of prototypical human cortical microcircuits with different levels of depression severity, in terms of reduced inhibition from SST interneurons (**Fig. 1A**). The models included four key neuron types (Pyr neurons and SST, PV and VIP interneurons) with type-specific synaptic properties and connection probabilities. Depression severity ranged from 0 to 40% reduced SST-mediated synaptic and tonic inhibition onto all cell types in the microcircuit. We simulated α5-PAM dose effects on the apical dendrites of Pyr neurons, ranging from 0 to 150% relative to the effect of a reference 3 μM dose of α5-PAM (ligand GL-II-73) on human cortical Pyr neurons, which we had characterized experimentally and modeled previously (see Methods). We simulated baseline (resting-state) activity in the microcircuits, together with dipole moments from which we calculated EEG signals. We thus simulated dose responses in 100 “virtual subjects” (microcircuits with randomized connectivity and activity) with different levels of depression severity (**Fig. 1B)**. EEG power increased with SST loss severity (**Fig. 2A**), in particular in the θ frequency band (4 - 8 Hz; Pearson correlation, *r* = 0.69, *p* = 1.52e-15; **Fig. 2B**) and α frequency band (8 - 12 Hz; *r* = 0.75, *p* = 1.90e-19; **Fig. 2C**), as well as in the broadband (3 - 30 Hz) range of the 1/*f* component (*r* = 0.71, *p* = 1.83e-16; **Fig. 2D**).

**Figure 1.**
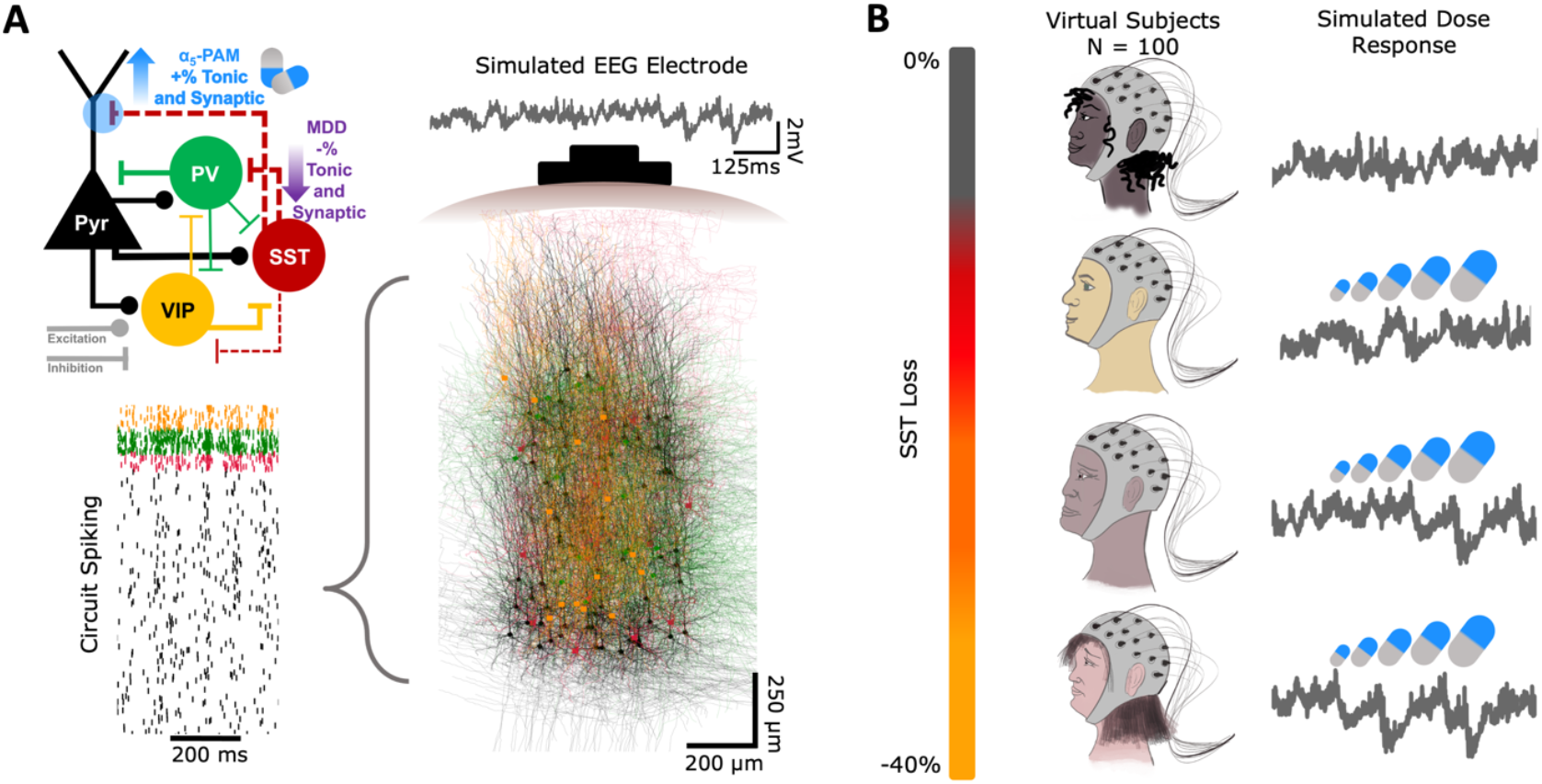
Simulating EEG of inhibition loss severity in depression and treatment response. **A**. Simulated neuronal spiking and EEG in human cortical L2/3 microcircuits in health, depression and under application of α5-PAM. Microcircuits were comprised of 1000 detailed neuron models of four types: Pyr (black), SST (red), PV (green), and VIP (yellow). The connectivity schematic (top left) highlights the cell-specific connectivity, the mechanisms of depression (MDD; loss of SST tonic and synaptic inhibition onto all cell types) and α5-PAM doses (boosted SST tonic and synaptic inhibition to Pyr neurons). **B**. We simulate five levels of SST inhibition loss severity (0%, 10% … 40%) across 20 different circuits each, representing a total of 100 different virtual subjects. For each subject we simulated a dose-response of α5-PAM (0%, 25%, 50% … 150% relative to the reference dose) and identified ground-truth optimal dose and range.

**Figure 2.**
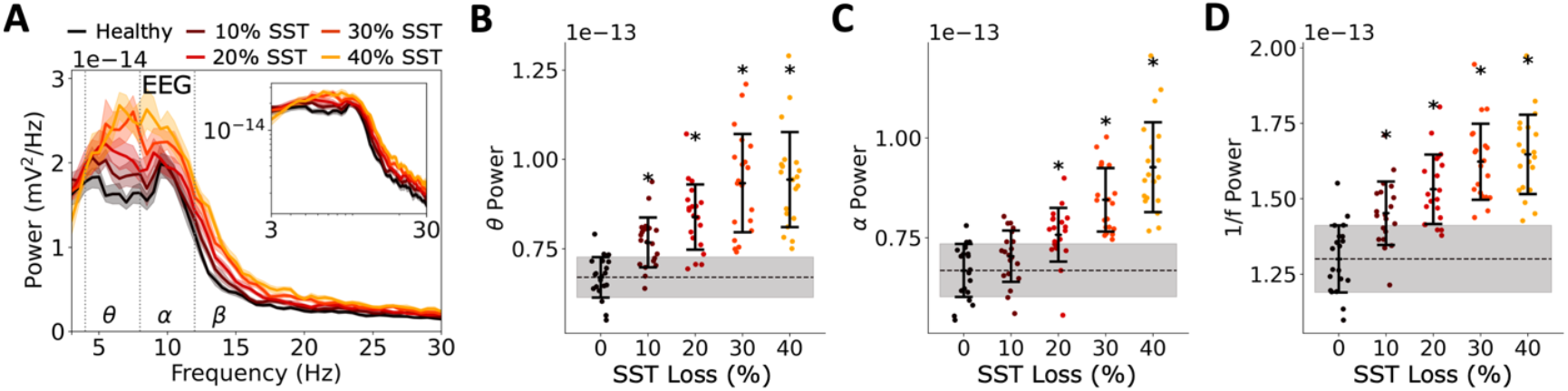
Simulated EEG biomarkers of depression severity. **A**. PSD of simulated EEG from each severity level of SST loss (bootstrapped mean and 95% confidence intervals across circuits). Inset – PSD plotted in log scale. **B-D**. PSD power in theta band (4 - 8 Hz, B), the alpha band (8 - 12 Hz, C), and broadband 1/*f* (3 - 30 Hz, D) for each level of SST loss (grey – healthy standard deviation; dashed line – healthy mean; error bars = mean and standard deviation). All asterisks denote significant paired t-tests (p < 0.05) with effect sizes greater than 1 when compared to healthy.

For each virtual subject, we computed the PSD following each α5-PAM dose administration (**Fig. 3A-B**) and extracted PSD metrics (1/*f*, α, and θ power). We fitted a dose-response function (relating dose and PSD metrics: Dose = m_1_·1/*f* + m_2_·θ + m_3_·α + b) and used it to identify the optimal dose and optimal dose ranges that would bring the subject’s EEG features to the healthy mean and the healthy ranges, respectively (**Fig. 3C**). We then used the optimal doses to fit a dose prediction model, using multivariate linear regression with the depression PSD metric values as input features (**Fig. 3D**). Prediction accuracy for the test subset of virtual depression subjects was high (90% ± 5%, n = 50 permutations of fit/test subject sets) and better than using univariate linear regression models with single PSD feature (1/*f* : 77% ± 6%, *p* = 3.25e-18, Cohen’s *d* = -2.1; θ: 84% ± 7%, *p* = 3.72e-6, Cohen’s *d* = -1.0; α: 79% ± 7%, *p* = 4.25e-15, Cohen’s *d* = - 1.9; **Fig. 3E**). For the cases where the dose prediction models were incorrect, they did not tend to underestimate or overestimate the correct dose (under-estimation: 4.7% ± 4.4%, over-estimation: 5.3% ± 2.8%, *p* > 0.05, Cohen’s *d* = 0.1). We selected the dose prediction model with the highest accuracy (predicted dose = 0.31·1/*f* + 0.77·θ + 1.18·α - 0.64) and used it for the rest of the analysis. The predicted doses recovered EEG features for the test subset of virtual depression subjects to the healthy range (normalized 1/*f* : 0.46 ± 0.11 vs 0.48 ± 0.13, *p* > 0.05, Cohen’s *d* = -0.2; normalized θ: 0.20 ± 0.10 vs 0.25 ± 0.10, *p* > 0.05, Cohen’s *d* = 0.5; normalized α: 0.39 ± 0.10 vs 0.34 ± 0.07, *p* > 0.05, Cohen’s *d* = -0.5; **Fig. 3F**). Inclusion of periodic features did not improve accuracy compared to using raw α and θ power and aperiodic 1/*f*.

**Figure 3.**
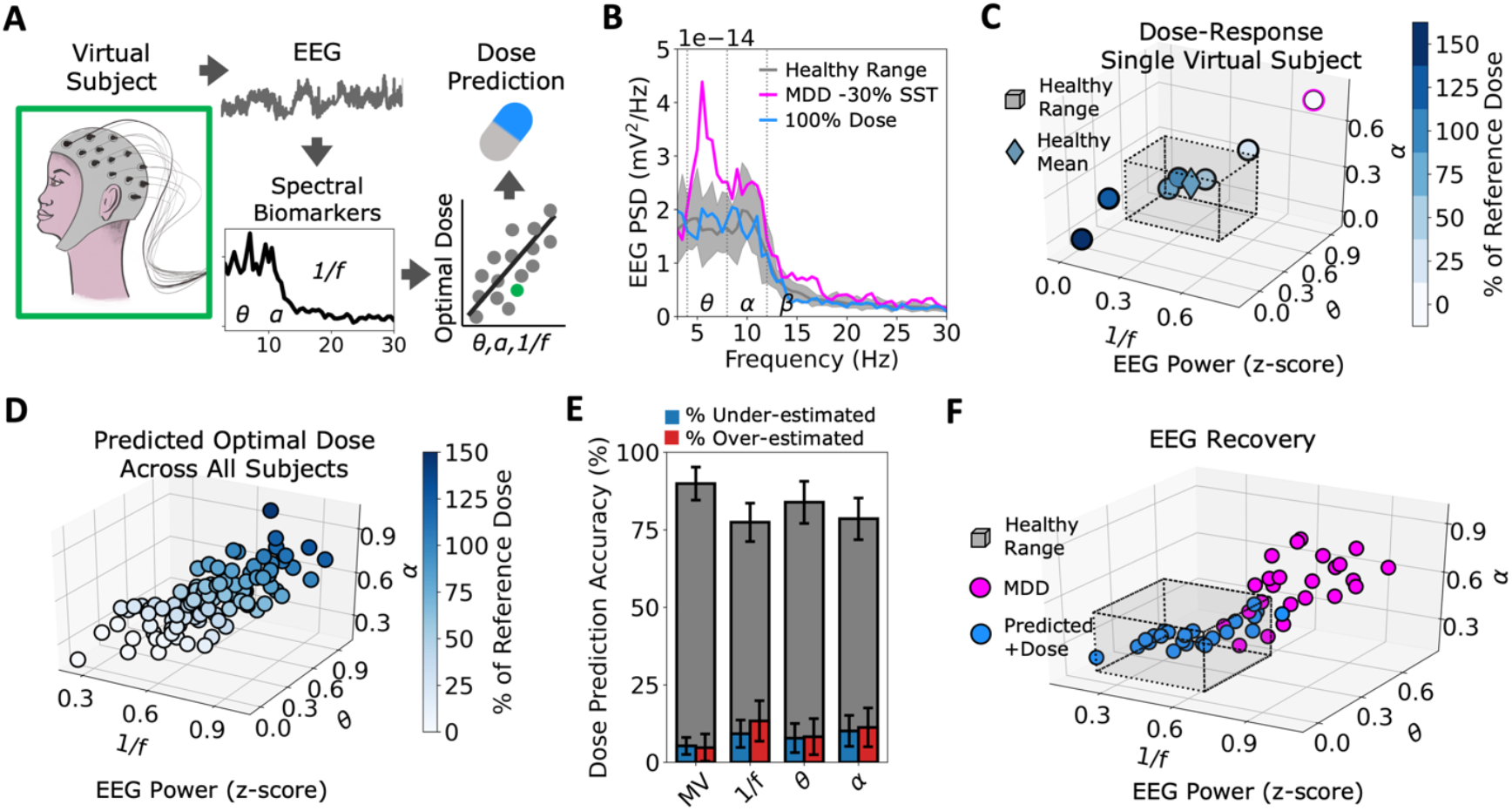
EEG biomarkers of depression severity predict α5-PAM dose accurately. **A**. Schematic illustrating the approach - we found optimal α5-PAM dose for each virtual subject based on power spectral biomarkers of their simulated EEG, and used the optimal doses to develop dose prediction models for restoring the EEG metrics back to healthy ranges across subjects. **B**. Example PSD profiles for one virtual subject (with 30% reduced SST interneuron inhibition, magenta), and under application of 100% of the reference α5-PAM dose (blue). Healthy mean and full ranges are shown in grey. **C**. Dose-response for the same virtual subject as in B, with 30% reduced SST interneuron inhibition (circle outlined in magenta), plotted across three EEG features (1/*f*, θ, α). A fit of the response was used to obtain the optimal dose (diamond color) and range with respect to the healthy EEG mean (diamond position) and ranges (grey cube), respectively. **D**. Predicted doses for each virtual subject (including healthy) as a function of its EEG features at baseline (before α5-PAM application). **E**. Percent of correct dose prediction for test sets of virtual subjects, for dose prediction models using either multivariate (MV) or single EEG biomarkers (50 permutations; blue = under-estimated errors; red = over-estimated errors). **F**. EEG metrics of all virtual subjects before (magenta) and after applying the predicted optimal α5-PAM dose (blue).

The model’s predicted dose based on EEG biomarkers also recovered microcircuit spiking and function in terms of failed and false detection rates, as measured from pre- and post-stimulus Pyr neuron spike rate distributions below and above the signal detection threshold, respectively. We compared the simulated baseline spiking activity for each virtual depression subject in the test subset and each dose to spiking activity following a brief stimulus (**Fig. 4A**) and calculated the proportion of failed and false detection errors based on microcircuit baseline and response spike rates (**Fig. 4B**). We then estimated the functional recovery due to the predicted dose from the dose-response curves for Pyr neuron spike rate (**Fig. 4C**), failed detections (**Fig. 4D**) and false detections (**Fig. 4E**), for each virtual depression subject in the test subset. We used a linear fit for the spike rate curves, and exponential fits for the detection rates curves. The predicted doses recovered the spike rates to the healthy ranges (0.76 ± 0.02 Hz vs 0.74 ± 0.07 Hz, *p* > 0.05, Cohen’s *d* = -0.4, **Fig. 4F**), as well as failed detection rates (1.50 ± 0.25% vs 1.32 ± 0.59%, *p* > 0.05, Cohen’s *d* = - 0.4) and false detection rates (1.59 ± 0.31% vs 1.42 ± 0.74%, *p* > 0.05, Cohen’s *d* = -0.3). For some virtual subjects with 10 - 30% SST interneuron inhibition loss, the predicted doses based on EEG biomarkers were slightly over-estimated, as measured by functional metrics, and for some virtual subjects with 40% reduced SST interneuron inhibition the predicted doses were slightly under-estimated, although they still brought the functional metrics much closer to the healthy range compared to before α5-PAM application.

**Figure 4.**
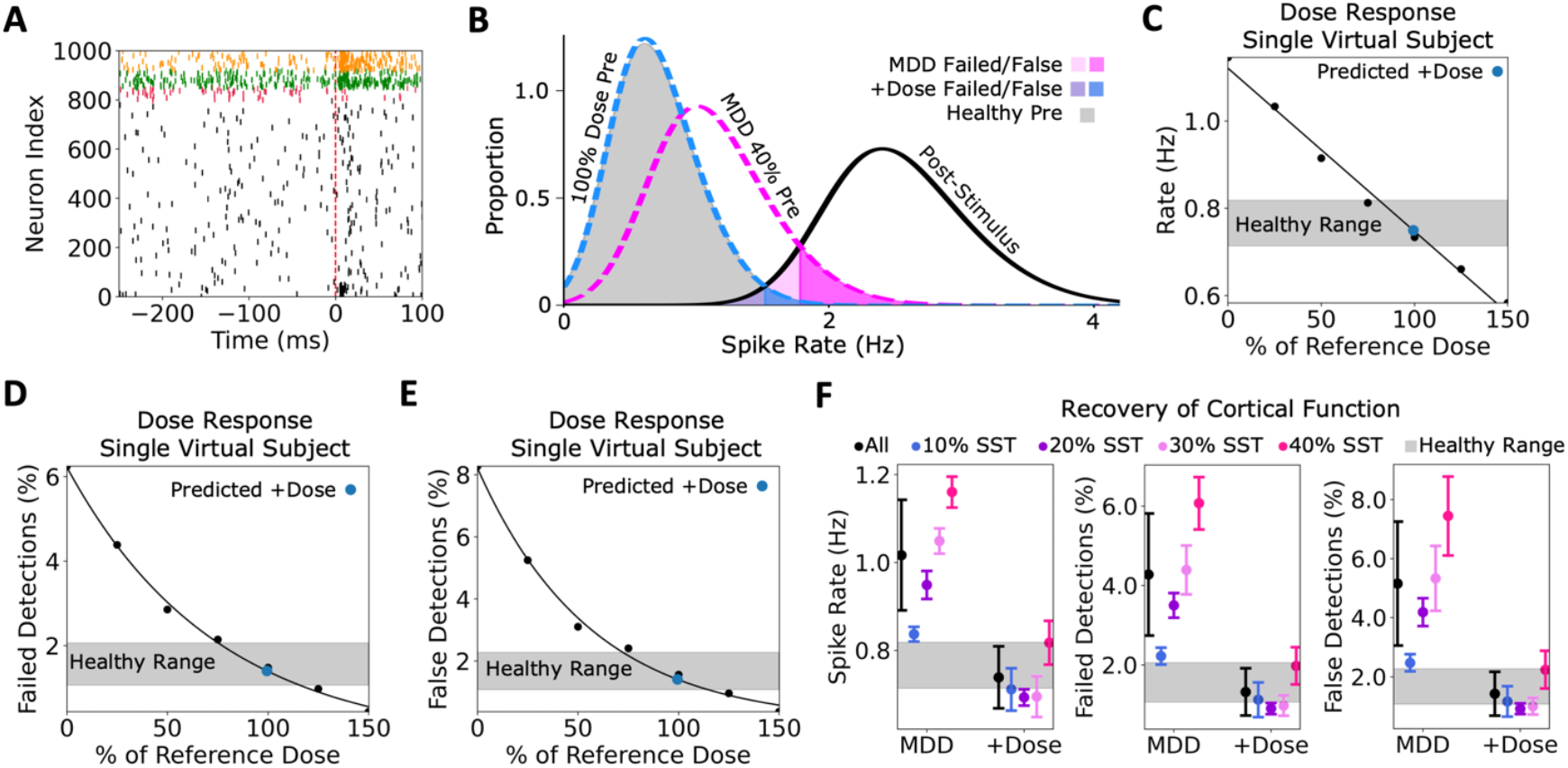
Predicted α5-PAM dose using EEG biomarkers recovers microcircuit spiking and function. **A**. Example raster plot of simulated baseline spiking and response to a brief stimulus. Dashed line indicates stimulus time. Cell type color code is the same as in figure 1A. **B**. Distributions of pre-stimulus firing rates from an example circuit with 40% SST inhibition reduction before (magenta) and after (blue) application of predicted dose. Average post-stimulus firing rates were similar across conditions, and are shown in black solid line. The overlaps between pre- and post-stimulus curves indicate failed and false signal detection errors. **C - E**. Dose-response curves in terms of Pyr neuron spike rate (C), failed detection rates (D) and false detection rates (E) for an example virtual subject with severity 40% SST inhibition reduction. The predicted dose based on EEG and the corresponding functional metrics is shown by the blue dot. **F**. Mean and standard deviation of spike rate (left), failed detection rates (middle), and false detection rates (right) in simulated depression (MDD) microcircuits before and after applying the predicted optimal dose. Grey area shows the healthy range.

Dose prediction models using an artificial neural network (ANN) or support vector machine (SVM) with multivariate EEG biomarker inputs had comparable dose prediction accuracy as using linear regression (ANN: 90% ± 3.5%, SVM: 92% ± 5.3%). Predicted doses using the ANN recovered EEG features (normalized 1/*f* : 0.49 ± 0.11, *p* > 0.05, Cohen’s *d* = 0.3; normalized θ: 0.25 ± 0.11, *p* > 0.05, Cohen’s *d* = 0.4; normalized α: 0.35 ± 0.10, *p* > 0.05, Cohen’s *d* = -0.3), spike rates (0.78 ± 0.09 Hz, *p* > 0.05, Cohen’s *d* = 0.3), as well as failed detection rates (1.68 ± 0.84%, *p* > 0.05, Cohen’s *d* = 0.3) and false detection rates (1.85 ± 1.06%, *p* > 0.05, Cohen’s *d* = 0.3). Predicted doses using SVM also recovered EEG features (normalized 1/*f* : 0.45 ± 0.13, *p* > 0.05, Cohen’s *d* = -0.09; normalized θ: 0.21 ± 0.10, *p* > 0.05, Cohen’s *d* = 0.2; normalized α: 0.34 ± 0.10, *p* > 0.05, Cohen’s *d* = -0.4), spike rates (0.73 ± 0.10 Hz, *p* > 0.05, Cohen’s *d* = -0.4), as well as failed detection rates (1.28 ± 0.74 %, *p* > 0.05, Cohen’s *d* = -0.4) and false detection rates (1.36 ± 0.88 %, *p* > 0.05, Cohen’s *d* = -0.3).

When generating dose prediction models that used Pyr neuron spike rates (Dose = 2.66·rate - 2.11) instead of EEG to predict doses, recovery of functional metrics was improved (**Fig. 5A**). Pyr neuron spike rates recovered to the healthy ranges for all virtual depression subjects in the test dataset (0.76 ± 0.01 Hz, *p* > 0.05, Cohen’s *d* = -0.06), as were the failed detection rates (1.40 ± 0.15%, *p* > 0.05, Cohen’s *d* = -0.5) and false detection rates (1.49 ± 0.18%, *p* > 0.05, Cohen’s *d* = -0.4). Dose prediction based on Pyr neuron spike rates also recovered the EEG features back within healthy ranges in 96% (22 of 23) of test subjects (**Fig. 5B)**, with group statistics not significantly different from healthy for 1/*f* (0.49 ± 0.09, *p* > 0.05, Cohen’s *d* = 0.3) or α (0.37 ± 0.06, *p* > 0.05, Cohen’s *d* = -0.1), although exhibiting a slightly larger θ power (0.26 ± 0.09, *p* = 0.049, Cohen’s *d* = 0.6) due to the one test subject that did not recover fully within healthy ranges.

**Figure 5.**
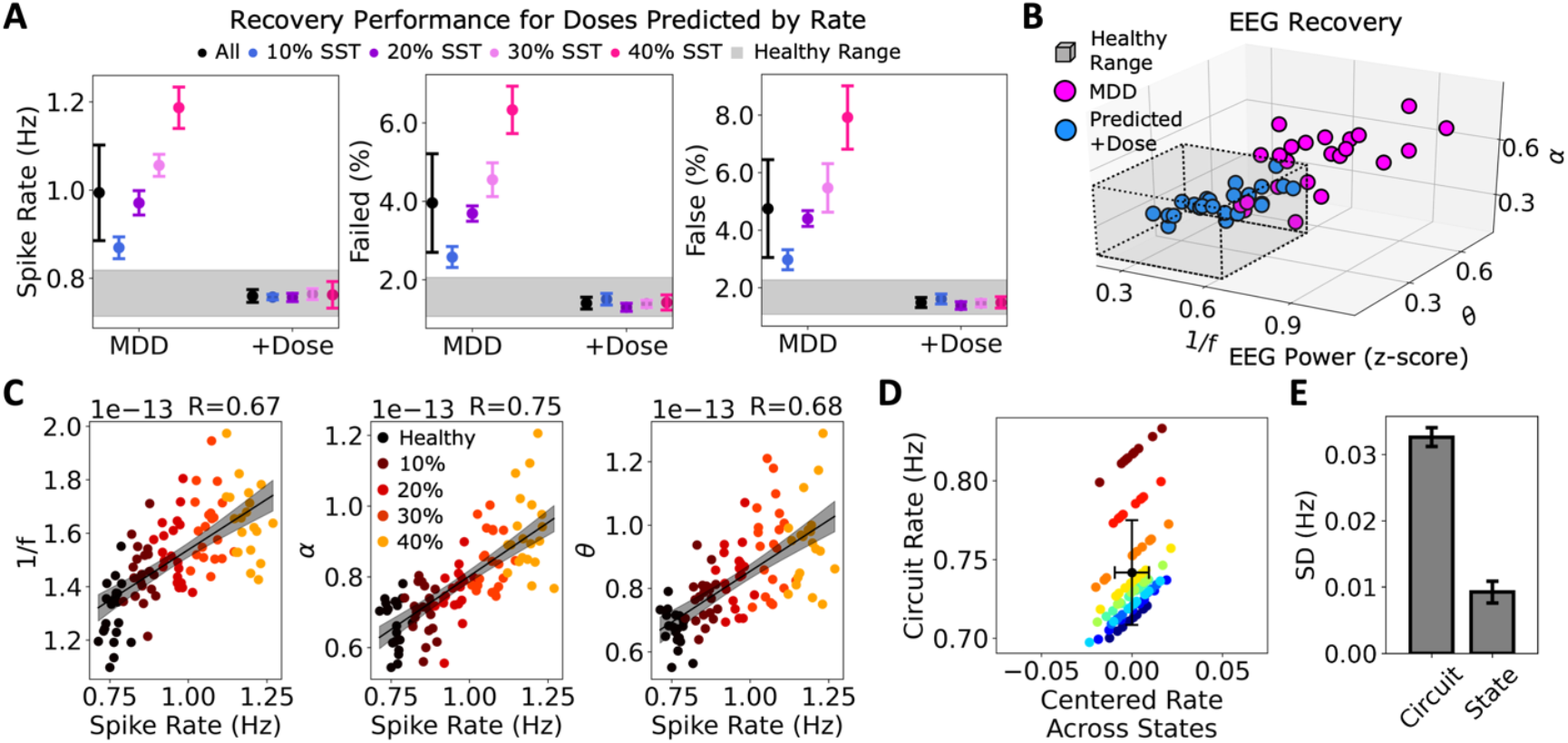
Dose prediction and recovery using microcircuit spike rates. **A**. Mean and SD of spike rates (left), failed detection rates (middle), and false detection rates (right) in virtual depression subjects (MDD) and after applying the predicted doses. Grey area shows the healthy range. **B**. EEG metrics of all virtual subjects before (magenta) and after applying the predicted α5-PAM dose based on spike rate (blue). **C**. EEG metric (left: 1/*f*, middle: α, right: θ) correlations with Pyr neuron spike rate across all virtual subjects (black: healthy; color indicates % reduced SST interneuron inhibition). R values are shown in the top right. **D**. Variance across virtual subjects was larger than the variance due to state (10 healthy microcircuits across 10 activity states). Error bars show SD. Different microcircuits are denoted by different colors. **E**. Average SD of spike rate across microcircuits or states.

Microcircuit spike rates therefore served better than EEG as biomarkers of SST interneuron inhibition loss, and although EEG metrics were correlated with microcircuit spike rates, some of the variance was independent (1/*f* : *r* = 0.67, *p* = 2.55e-14; α: *r* = 0.75, *p* = 4.56e-19; θ: *r* = 0.68, *p* = 9.94e-15, **Fig. 5C**). Interestingly, baseline spike rate variability was driven largely by differences between virtual subjects (microcircuit synaptic connectivity) rather than between activity states (the timing of background inputs to the microcircuits) within subjects (variance across healthy virtual subjects: 0.033 ± 0.001 Hz, variance across states: 0.009 ± 0.002 Hz, *p* = 2.14e-17, Cohen’s *d*: -14.4; **Fig. 5D-E**). Therefore, differences in microcircuit connectivity between virtual subjects resulted in some subjects tending to have hypo-active baseline firing and others hyper-active firing, which consequently influenced the required dose. Virtual subjects were also distinguished as being either hypo- or hyper-active in terms of their EEG metrics, although there was no significant difference in the variance across virtual subjects compared to activity states (1/*f*: *p* > 0.05, Cohen’s *d*: -0.4; α: *p* > 0.05, Cohen’s *d*: 0.02; θ: *p* > 0.05, Cohen’s *d*: -0.2).

## Discussion

In this work, we showed *in silico* that α5-PAM therapeutic doses, which effectively recover cortical microcircuit EEG, spiking activity and function, can be predicted from simulated EEG biomarkers of depression severity in terms of reduced SST interneuron inhibition. We used detailed simulations of human microcircuits to mechanistically link severity of SST interneuron inhibition loss, EEG biomarkers and effective α5-PAM doses. Our machine learning predictors and candidate EEG biomarkers could serve to facilitate translation efforts of α5-PAM pharmacological treatment by stratifying depression patients that may benefit from α5-PAM, and by predicting treatment outcome and monitoring α5-PAM efficacy with non-invasive brain signals.

We modelled depression severity in terms of SST interneuron inhibition reduction, and thus the tools we developed would be relevant to a subset of depression patients, which could amount to 50% of the patients as indicated postmortem^16^. Some of the relevant EEG biomarkers we found, such as power in the theta and alpha frequency bands, have also played a role in previous machine learning methods that stratified depression patients by EEG features^35–37,41^. However, the “black box” machine learning methods are generally blind to any underlying mechanisms and are therefore limited in terms of accurately stratifying a particular mechanistic subtype^41^. Most previous machine learning studies also focused on classifying depression patients from healthy controls^41,47^ rather than estimating severity. In addition, there have been only a few studies that found correlations between EEG features and severity of depression symptoms, which were limited to higher (beta and gamma) frequency bands^48,49^ or theta cordance following antidepressant treatment^50^. More work is therefore needed to validate the severity biomarkers we have characterized.

We constrained the models with the effect of 3 μM of the α5-PAM compound, GL-II-73 (i.e., the reference 100% dose) as previously^11^, which is in the range for selectively targeting α5 subunit receptors without substantially activating other subunits^9,10^. In rodents, this dose was also within the range that yielded anxiolytic, antidepressant, and pro-cognitive effects without sedation^10^. To maintain the interpretability of our models in only selectively targeting of α5-GABA_A_ receptors, we limited the upper range of doses we simulated to 150% of the reference dose, since higher doses are expected to have broader effects of boosting inhibition non-selectively^9,10,51^. In some virtual subjects with severe SST reductions the upper-range dose was necessary for recovery, but for most virtual subjects the required doses were lower, supporting the dose range we examined.

We used our previous detailed models of human cortical L2/3 microcircuits that capture the effects of depression mechanisms on cortical microcircuit function and human resting state EEG signals^11,28,30^. Future studies could refine our results and methods by including deeper layer circuits^52–54^, layer 1 interneurons^55,56^, and additional mechanisms of depression^9,57^. The machine learning models for dose prediction we have developed required only a few key EEG features and we selected these features based on whether they exhibited strong correlations with reduction in SST interneuron inhibition. Future implementations could consider alternative and more systematic feature selection methods for linear regression or SVM models, and alternative network architectures and feature sets for the ANN models. Nevertheless, the different machine learning models predicted dose with high accuracy and performed comparably well, indicating that the features we chose were sufficient.

In this work we provide the first demonstration of α5-PAM dose prediction using simulated EEG biomarkers of reduced SST interneuron inhibition severity in detailed depression microcircuit models with α5-PAM effects. Our study overcomes limitations of doing the above in living humans, and thus our tools could serve to better stratify depression patients that may benefit from α5-PAM treatment and enable EEG-based treatment outcome prediction and monitoring efficacy.

## Methods

### Models of human cortical microcircuits in health and depression

We used morphologically- and biophysically-detailed models of human L2/3 cortical microcircuits in health and depression, as described previously^11,28^. The models were comprised of 1000 neurons (80% Pyr, 5% SST, 7% PV, and 8% VIP) distributed across a 500x500x950 μm^3^ volume, and reproduced neuronal firing and synaptic properties as measured in human neurons. For complete list of data provenance in our models, please refer to supplemental tables in our previous work^28^. The models were simulated using NEURON 7.7^58^ and LFPy 2.0.2 (Python 3.7.6)^59^ on SciNet parallel computing^60^.

We modelled virtual healthy and depression subjects with different severity (0, 10, … 40%, n = 20 per group, n = 100 in total) in terms of reduced SST interneuron synaptic and tonic inhibition conductance onto all cell types^28^. Virtual subjects differed in microcircuit synaptic connectivity, background inputs, and spatial placement of neurons in the L2/3 volume.

### Simulating α5-PAM application

We simulated a reference dose of α5-PAM application corresponding to experimentally-measured effects of a 3 μM dose of GL-II-73 on human Pyr neurons, modelled as a 60% boost of synaptic and tonic inhibition to Pyr neuron apical dendrites^11^. We simulated the application of different doses of α5-PAM relative to this reference dose (25% to 150%, where 100% is the reference dose).

### Microcircuit baseline spiking activity

We simulated baseline activity in the microcircuit as described previously^11^, driving the microcircuit with background excitatory inputs of Ornstein Uhlenbeck (OU) point processes^61^. Independent excitatory OU point processes were placed at the midway points along the length of each dendritic arbor, and for Pyr neuron models, we placed 5 additional OU processes along the apical dendrites (at 10%, 30%, 50%, 70%, 90% of the apical length). The mean and standard deviation of each OU conductance were scaled up exponentially with relative distance from soma (ranging from 0 to 1) to normalize their effect.

### Simulated microcircuit EEG and power spectral analysis

Along with the baseline activity, we simulated resting-state EEG from the microcircuit models in LFPy 2.0.2 (Python 3.7.6) using a four-sphere volume conductor model (representing grey matter, cerebrospinal fluid, skull, and scalp with radii of 79 mm, 80 mm, 85 mm, and 90 mm, respectively) that assumed homogeneous, isotropic, and linear (frequency-independent) conductivity^11,30^. The conductivity for each sphere was 0.047 S m^−1^, 1.71 S m^−1^, 0.02 S m^−1^, and 0.41 S m^−1^, respectively^11,30,62^. We computed EEG power spectral density (PSD) using Welch’s method^63^ from the Scipy python module with 2s time windows. We also decomposed the EEG power spectra (in the 3–30 Hz range) into periodic and aperiodic components using the FOOOF toolbox^64^. The aperiodic component was a 1/f function parameterized by vertical offset and exponent parameters. We fitted the periodic oscillatory component with up to 3 Gaussian peaks defined by center frequency, bandwidth (min: 2 Hz, max: 6 Hz), and power magnitude (relative peak threshold: 2, minimum peak height: 0)^11,30^. We extracted EEG features including the area under the curve of power in different frequency bands (θ = 4 - 8 Hz, α = 8 - 12 Hz) and the 1/*f* aperiodic component (3 - 30 Hz range). Feature values were normalized by transforming to log10 space, z-scored relative to all conditions, and scaled to values ranging from -1 to 1.

### Optimal dose and optimal dose range estimations

For each virtual subject, we fitted a dose-response function (relating dose and EEG features), and identified the optimal dose and optimal dose ranges for the subject by finding the dose for which the function intersected with the EEG feature values of the healthy mean and the healthy ranges, respectively.

### Dose predictor models

We used the optimal dose and EEG features (1/*f*, θ and α power) of each of 70% of the subjects to fit a multivariate linear prediction model of the form:

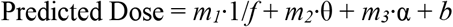

We tested the predictor performance using the remaining 30% of subjects, in terms of the proportion of correct dose predictions. A dose was correct if it fell within the subject’s optimal dose range. We generated multiple models using 50 train/test permutations, and selected the best performing prediction model.

As an alternative type of model, we generated predictor models similarly but instead used a linear support vector machine (SVM) in python^65^, with a linear kernel, a regularization parameter of 10, a tolerance value of 0.001, and an epsilon value of 0.1. As another type of predictor model, we trained an artificial neural network (ANN) using the tensorflow python package^66^. The ANN comprised of 3 input nodes (corresponding to the EEG features above), 9 hidden layer nodes with ReLU activation, and 1 output layer node with linear activation. For learning, we used an Adam optimizer with a learning rate of 0.01, mean absolute error as the loss and training accuracy functions, and initialized weights from a normal distribution centered at zero. In both SVM and ANN cases, we fit the models using 70% of the data and tested using 30%. For the SVM we ran 50 train/test permutations, and for the ANN we ran 10 train/test permutations (for computational efficiency), from which we selected the best performing prediction model.

### Failed/false detection metrics

We calculated failed and false detection errors in the different conditions using the distribution of Pyr neuron firing rates at baseline (n = 23,950 windows of 50 ms for each microcircuit) and a reference distribution of simulated firing rates in response to brief stimulus (calculated across 200 stimulus presentation, in the 5-55 ms period post-stimulus), since we have previously shown that response rate was not impacted by SST loss alone or application of α5-PAM^11,28^. The intersection point of the two distributions defined the stimulus detection threshold. We computed probability of false detections as the integral of the pre-stimulus distribution above the detection threshold divided by the integral of the entire pre-stimulus distribution. Similarly, we computed the probability of failed detections as the integral of the post-stimulus distribution under the detection threshold divided by the integral of the entire post-stimulus distribution.

### Intra-versus inter-microcircuit variability analysis

To analyze the variance in spike rates due to state randomizations versus microcircuit randomizations, we simulated 10 randomized states across 10 randomized microcircuits (i.e., virtual subjects). Randomizing the microcircuit comprised of resampling synaptic connections and neuron positions in space, whereas randomizing the state comprised of resampling background OU noise inputs.

### Statistics

For linear correlations, we used two-sided Pearson correlations. For group comparisons we used two-sided paired- and independent-sample t-tests, where indicated. For independent-sample t-test cases where variance between groups were significantly different from each other (using the Levene test for equal variances) we performed a Welch’s t-test. Cohen’s *d* was calculated as follows:

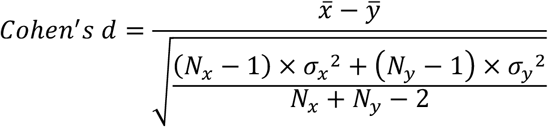

## Code availability

All original code has been deposited on Github and will be publicly available as of the date of publication.

## Acknowledgements

AGM and EH thank the Krembil Foundation for their generous funding support. AGM thanks the Labatt Family Network for Research on the Biology of Depression for funding support. Special thanks to A. Sherrington for contributing the face drawing in Fig. 1B.

## Author Contributions

Conception: AGM, TDP, ES, EH. Design: AGM, TDP, ES, EH. Analysis of the data: AGM, EH. Interpretation of the data: AGM, FM, TDP, ES, EH. Manuscript drafting: AGM, EH. Manuscript revisions: AGM, TDP, ES, EH. Manuscript final approval: AGM, TDP, ES, EH. Work accountability: AGM, EH.

